# A Phenomenological Model of Synaptic Activity in the Blowfly Lamina

**DOI:** 10.64898/2026.11.01.763551

**Authors:** Golam M Kashef, Rob de Ruyter van Steveninck

## Abstract

Estimates of the effective Poisson rate [1] obtained from voltage recordings of Large Monopolar Cells (LMCs) in the blowfly lamina, measured in response to repeated light-contrast stimuli, are on the order of 5 ×10^5^ events per second [2–4]. These values imply that if vesicle exocytosis at photore-ceptor–LMC synapses consisted of statistically independent quantal release events, the mean release rate would need to be at least this large to account for the observed signal and noise characteristics [2, 3]. Such a scenario is highly implausible, as it would correspond to an exocytosis rate of approximately 400 vesicles per second per active zone, or equivalently to the turnover of a membrane area comparable to that of all six photoreceptor axon terminals within a lamina cartridge more than twice per second [2, 3]. Relaxing the assumption of statistical independence, however, allows for substantially larger effective Poisson rates. In earlier work [2, 3], we demonstrated that temporal regularization of release events—particularly through a reduction in low-frequency noise power spectral density—can increase the effective Poisson rate without altering the underlying mean release rate. Additionally, the contrast-to-current impulse response function of the extracellular space has been shown to be approximately equal in magnitude and opposite in sign to that of the LMCs [2, 3], suggesting that both signals arise from the same chloride current. This observation provides a potential physiological mechanism for vesicle regularization, although it was previously described only in qualitative and simplified terms. In the present work, we develop a phenomenological model that incorporates these findings together with additional physiological parameters from literature to account for the regularization of vesicle release.

---

Adult flies, like most adult insects, possess compound eyes. In the blowfly, each eye comprises roughly 5,000 ommatidia, each containing photoreceptors that form the outer retina. These photoreceptors transduce incident photons into electrical signals and relay their outputs synaptically to Large Monopolar Cells (LMCs) within the lamina, the first optic neuropil. Despite carrying out what appears to be a relatively simple transformation of photoreceptor signals, the lamina represents a major investment of neural resources. In the blowfly, it constitutes approximately 10% of total brain volume, contains about 20% of all neurons, and consumes roughly 2% of resting metabolic power. This disproportionate investment reflects the behavioral demands of rapid, agile flight, which require visual processing that is both fast and highly sensitive to reliably represent the environment during swift motion and abrupt turns [5].

The lamina is organized into repeating modular units called cartridges, analogous to the subdivision of the retina into ommatidia. Each cartridge contains the axon terminals of six photoreceptors (R1–R6) originating from six different ommatidia but sampling the same direction in visual space through neural superposition [6, 7]. As a result, convergence effectively increases signal reliability by reducing noise without loss of spatial information. Each cartridge is a cylindrical, glia-ensheathed structure containing 14 identified cell types. Two nearly identical LMCs, L1 and L2, are the primary postsynaptic targets of photoreceptors, each receiving input from six photoreceptors and accounting for the majority of photoreceptor synapses. Unless otherwise stated, references to LMCs hereafter denote these two cells.

Photoreceptor terminals contain on the order of 220 active zones each, organized as tetrad synapses in which a single presynaptic site contacts four postsynaptic elements, including both L1 and L2 [8]. Individual cartridges are relatively electrically isolated, and light-driven activity generates substantial extracellular potentials [2, 9]. Under repeated contrast stimulation, both LMCs and the extracellular compartment exhibit similarly high effective Poisson rates (derived from the ratio of the contrast power transfer and noise power spectral density [1, 2]. These measured rates are substantially higher than the rate for a single photoreceptor, though they generally do not quite reach the theoretical maximum of six times the rate of a single photoreceptor[2]. The LMC voltage response arises from a fundamentally discrete process-the release of synaptic vesicles from photoreceptor active zones. Vesicle release is initiated by the opening of voltage-gated calcium channels in the active-zone membrane, allowing Ca^2+^ ions to enter the photoreceptor terminal and trigger a biochemical cascade that culminates in vesicle exocytosis. In general, a higher photoreceptor transmembrane voltage increases the probability of channel opening and therefore elevates the rate of vesicle release. Each exocytotic event releases histamine into the extracellular space, which activates ligand-gated chloride channels in the LMC membrane, permitting Cl^−^influx into the LMC. The resulting inward chloride current drives the LMC membrane potential in the negative direction, accounting for the sign-inverting nature of the synapse. If we were to go with the assumption that individual events are statistically independent (i.e. occurring through a Poissonian process), the measured effective Poisson rate suggests an average rate of at least 5 × 10^5^ events per second, which is a rather unfeasible given spatial constraints. [2]

Previous work using repeated light contrast stimulation and injected current has shown that the contrast-to-current impulse response function of the extracellular compartment (computed from contrast-to-voltage and current-to-voltage impulse responses) is approximately equal in magnitude and opposite in sign to that of an LMC. Setting aside the caveat that the extracellular current should, strictly speaking, reflect the combined contributions of two LMCs rather than one, this observation suggests that current leaving the extracellular space during visual stimulation is effectively the same current that enters an LMC [2]. This framework allows a single vesicle release event to be viewed not only as transiently hyperpolarizing the LMC via chloride influx, but also as increasing (i.e. depolarizing) the extracellular potential. The transient increase in extracellular potential will reduce the photoreceptor transmembrane voltage and suppress further vesicle release, until relaxation restores the release rate. The resulting time-dependent correlations imply that vesicle release events are no longer statistically independent. Regularization of release events could substantially increase the effective Poisson rate relative to the true event rate by reducing low-frequency noise. This is tied to the fact that the predictability of regularized arrivals makes it possible to infer the modulated rate (at least at the frequencies that fall below the average rate of release) from fewer samples.

The computational model presented here explicitly incorporates the dependence of vesicle release rate on the photoreceptor transmembrane potential, generating a stochastic sequence of release events that drive chloride current into the LMC. The current-to-voltage impulse response function reported by Kashef and de Ruyter (2026) is then used to map this chloride current onto voltage responses in both the LMC and the extracellular compartment. This construction closes a feedback loop between synaptic release and the photoreceptor transmembrane voltage, providing a mechanistic substrate for the emergence of vesicle release regularization.

## I. METHODS

Given that it is a feedback system where the current state of the output is fed back as input to generate future states, and that there are multiple components working in tandem to produce output, it is difficult to explain our computational model in a linear fashion. Rather, we first refer the reader to Figure 1, which presents all the variables in this simulation and the operations connecting them as a flowchart. Below, we discuss each variable one by one, explaining how they are implemented, the relationships they have to other variables, and how those relationships are implemented. A time step in our simulation represents *δt*_sim_ = 2µs,or, in other words *f*_sample,sim_ = 500kHz. The model incorporates experimental data that was collected or processed at *f*_sample,exp_ = 5kHz. Wherever we encounter an experimentally derived input (shown with blue dotted boxes in Figure 1), the imported data is thus upsampled from 5 kHz to 500 kHz using the Piecewise Cubic Hermite Interpolating Polynomial (Pchip) method. This method adds smoothness while being shape-preserving, maintaining the original monotonicity when going from one original sample to the next[10].

**FIG. 1.**
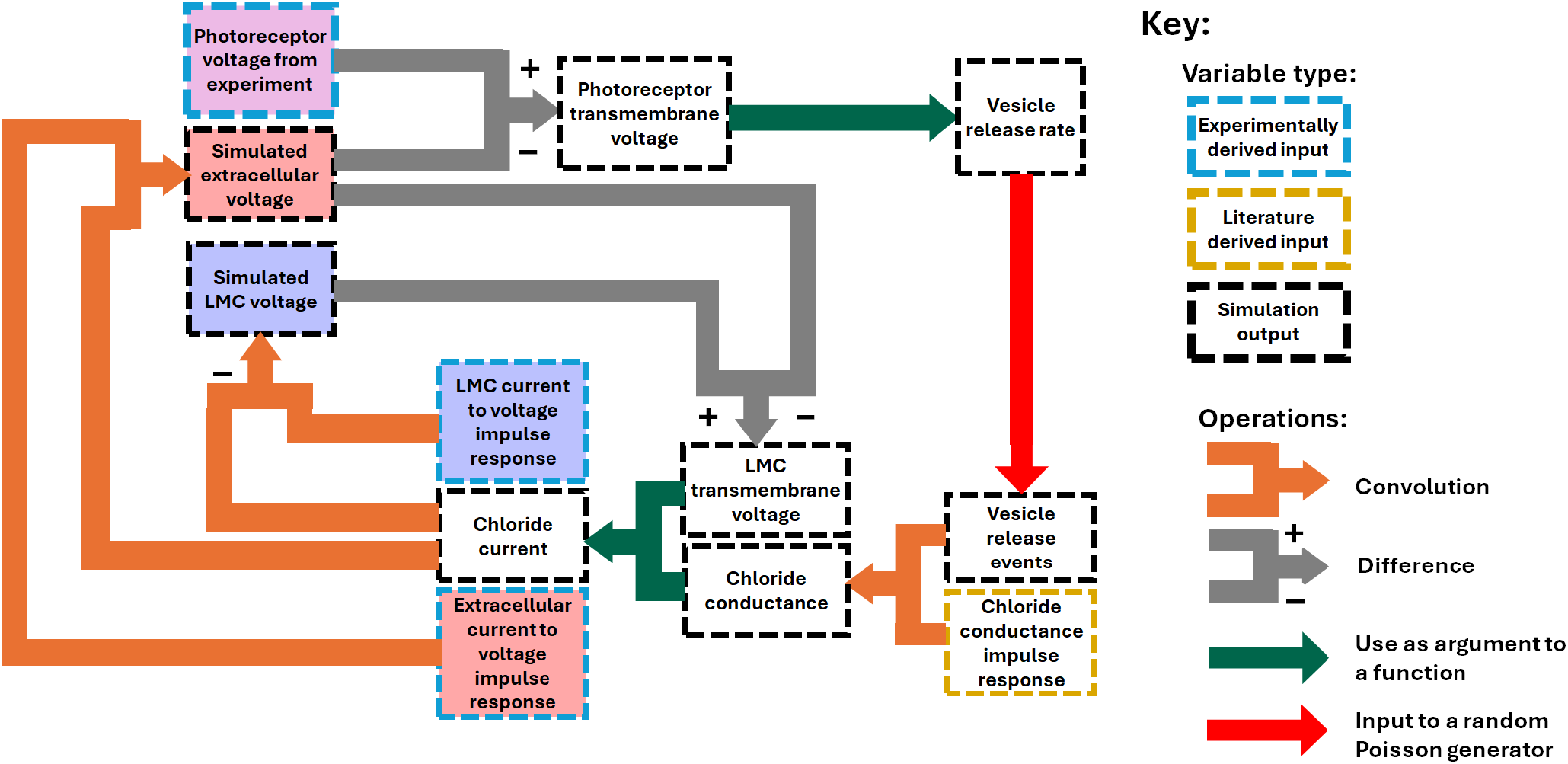
Flowchart representing the variables and the operations connecting them in our numerical simulation. A key is provided on the right, explaining the source/role represented by the border color of each variable, and the type of operations that are connecting them. The negative sign on one of the convolutions represents sign inversion.

### A. Photoreceptor voltages

200 photoreceptor voltage traces stimulated by a repeated 500ms light contrast in a previous study [2] were taken as one long time series, and 200 copies of the ensemble averaged signal were stitched together and subtracted from this, to give us a long “noise” trace. The light contrast function used to stimulate the photoreceptor has a flat spectrum and a standard deviation of *σ*_*c*_ = 0.25. Since six photoreceptors feed to an actual lamina cartridge and process information in parallel, we may replace the effect of the six photoreceptor voltages with a single simulated photoreceptor voltage *V*_PR,sim_ given by the average voltage of the six photoreceptors. As each photoreceptor in a single LMC collects light from a single direction in space, we may think of them as having the same signal, and only varying by noise. Calling the signal *V*_PR_(*t*) and the noise *η*_PR*i*_(*t*) for each photoreceptor voltage *V*_PR,*i*_, the above equation may be written as

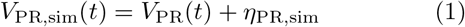

where

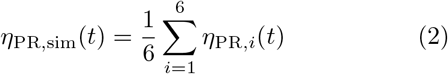

In practice, we create a set of six different noise traces by circularly shifting the noise time series by each of the first six integer multiples of ⌊200*/*6⌋ = 33 trial lengths. Then the input to the simulation is *V*_PR,sim_(*t*) as defined in Equation 1 (see top panel in Figure 4).

### B. Instantaneous release rate from Δ*V*_PR_

At a photoreceptor active zone, a greater transmembrane voltage Δ*V*_PR_ opens up more voltage gated Ca^2+^ channels, and the incoming Ca^2+^ acts as a catalyst for vesicle exocytosis. Ignoring some microscopic details, we may think of a single Ca^2+^ channel as a two-state system-it is either open or closed. Let the natural energy of a single channel when it is open be *ε*_1_ and when it is closed be *ε*_0_. Having a channel open lowers its energy by *ze*Δ*V*_PR_ where *z* is the number of charges on the voltage-gating molecule, and *e* is the elementary charge. Thus the energy of a single channel, *E*_*i*_, is given by

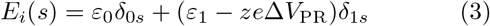

where *s* = 0 if the channel is open and *s* = 1 if the channel is closed. The Boltzmann probability for the channel to be open at any given time is given by

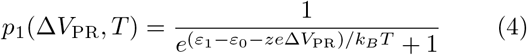

Defining the “activation voltage” as *V*_act_ = (*ε*_1_ −*ε*_0_)*/ze* and a logistic “slope” parameter *b* as *ze/k*_*B*_*T*, we continue to write the probability of a channel being open as

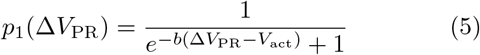

We may think of *p*_1_(Δ*V*_PR_) as the fraction of channels that are open at any given time. Then, defining *R*_max_ as the number of channels available times the rate of exocytosis events per open channel (i.e., the maximum possible vesicle release rate), the local time vesicle release rate is given by

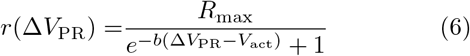

Equation 6 is the function of the instantaneous transmembrane voltage we will be using to define the instantaneous vesicle release rate in our simulations (see the relevant teal arrow in Figure 1).

We keep *R*_max_ and *V*_act_ as free parameters. Laughlin et al.(1987) show that single LMC signal amplitudes are related to single photoreceptor signal amplitudes by a logistic curve with a slope parameter of 0.56mV^−1^[11]. A similar slope parameter is observed in a logistic fit of a projected chloride conductance against the photoreceptor transmembrane voltage in Kashef and de Ruyter (2026) [2]. Since the LMC signal is driven by the vesicle rate, we assume that the shape of the saturating behavior car-ries over. We thus use *b* = 0.56mV^−1^ . Going back to the relation *b* = *ze/k*_*B*_*T*, we get a value of *z* = 14.4 at room temperature. Voltage-gated Ca^2+^ channels have four voltage-sensing domains (VSDs), each containing a positively charged S4 helix that moves in response to changes in the membrane potential. The S4 helix contains positively charged gating charges. While the exact number of gating charges varies between different VSDs, it is generally accepted that there are usually four gating charges per VSD [12]. This means that there are approximately 16 gating charges per channel, and our derived value of *z* = 14.4 is pretty close to this, giving us further reason to fix *b* at *b* = 0.56mV^−1^.

Simulating probabilistic release of synaptic vesicles involves calculating *r*(Δ*V*_PR_) at each time step, and using it to find the expected number of events at the time step as *µ*_*t*_ = *r*(Δ*V*_PR_) · *δt*_sim_. If *µ*_*t*_ *>* 0.25 we draw a number from a Poisson distribution with mean *µ*_*t*_, otherwise we do a Bernoulli trial to speed up the simulation, incurring a tiny cost in accuracy. Since the system is sensitive to single events, our tiny value of *δt*_sim_ = 2µs is chosen to make sure that we virtually encounter only 1 or 0 as the number of events in a time step. The random number generated is thus the number of vesicles released at that timestep.

### C. Event to Chloride Conductance Impulse Response and the Chloride Conductance

Here, we use the *in vitro* experimental results from Skingsley, Laughlin and Hardie (1995) [13], where LMCs were isolated from the lamina of various species of flies and inside-out patch recordings were performed on dendritic membranes containing histamine-activated Cl^−1^ channels. For the *C. vicina* LMC, the membrane was exposed to 10µM histamine, and spontaneous channel openings were observed at +50mV and −50mV patch voltages. The channels had variable periods of staying open, but displayed a less variable level of channel conductance when they opened. For *C. vicina* in −50mV, the mean conductance amplitude was found to be *G*_Cl_ = 60pS on average. The channel opening period histograms consistently suggest an exponential distribution, with the time constant found to be *τ*_open_ = 0.5ms for *C. vicina*.

Let us say that a single event opens up the number *N*_Cl_ of independent Cl^−^ channels. Then, if *N*_Cl_ is large, we can approximate the event delta function train to Cl^−^ conductance impulse response to be

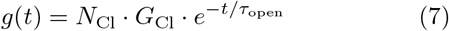

This allows us to define the time evolving Cl^−^ conductance in the LMC membrane as

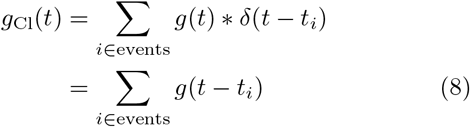

### D. The LMC transmembrane voltage and the synaptic chloride current

The LMC transmembrane voltage Δ*V*_*LMC*_ at a given time-step is found by taking the difference between the simulated LMC potential and the simulated extracellular potential at the time-step. Figure 2 shows the chloride conductance as a circuit element separating the LMC intracellular environment and the cartridge extracellular space, with the chloride reversal potential giving rise to an effective synaptic “battery”. From here, simply applying Ohm’s law gives us

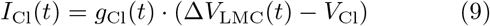

where *I*_Cl_(*t*), the synaptic chloride current, is the conventional current into the cartridge extracellular space from the LMC.

**FIG. 2.**
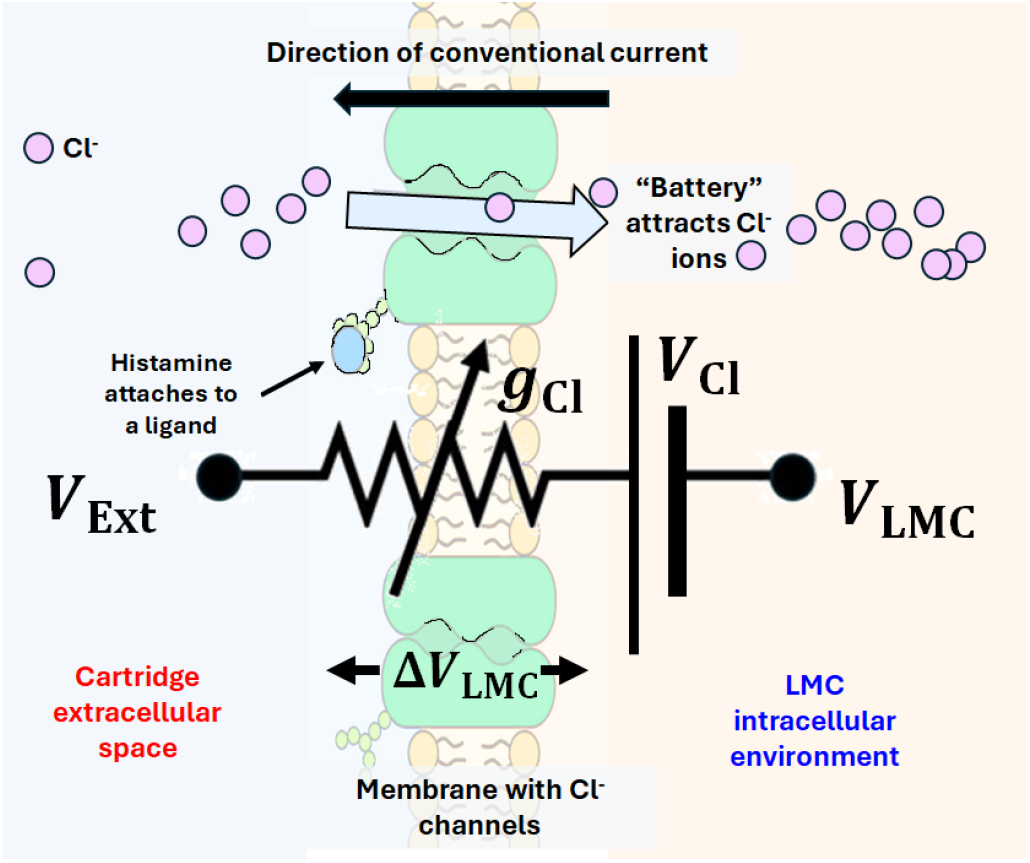
A combination of concentration and electrical gradients across an LMC membrane cause Cl^*−*^ ions to rush in when histamine causes a Cl^*−*^ to open. The attraction of the negatively charged Cl^*−*^ ions in the direction of the LMCs down the concentration gradient acts as a synaptic “battery”, whose voltage is called the Cl^*−*^ reversal potential *V*_Cl_. The opening and closing of channels as a result of histamine activity (i.e. vesicle exocytosis) is modeled as a time-varying chloride conductance *g*_Cl_. By Ohm’s law, the chloride current is thus given as *I*_Cl_ = *g*_Cl_(Δ*V*_LMC_ *− V*_Cl_)

The typical value of internal chloride ion concentration is 108mM, while that of the concentration outside is 560mM.[14, 15]. Plugging these into the Nernst equation for a temperature of 25^°^C, we get the following chloride reversal potential.

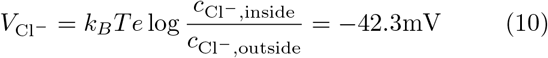

This reversal potential is corroborated by voltage clamp experiments by Laughlin and Osorio (1989)- the polarity of LMC light response switches when, through the application of constant current, the LMC is clamped to around 30mV to 45mV below its resting voltage [16].

### E. LMC and extracellular potentials from *I*_**Cl**_(*t*),*z*_**LMC**_(*t*) **and** *z*_**Ext**_(*t*)

Defining the synaptic chloride current, *I*_Cl_(*t*), allows us to define the LMC and extracellular potentials as

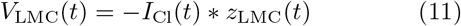

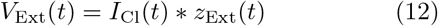

where *z*_LMC_(*t*) and *z*_Ext_(*t*) are specifically the injected current-to-voltage impulse response functions measured experimentally in Kashef and de Ruyter (2026)[2] (see Figure 7 of the work) by injecting random nanocurrent signals into the LMC and extracellular space respectively and using cross-correlation of the injected current and the voltage response. The negative sign in Equation 11 is from the fact that *I*_Cl_ is the conventional current moving *out* of the LMC.

The experiments in Kashef and de Ruyter (2026) also involved simultaneous stimulation of the early visual system by a repeated flat spectrum light contrast signal, to yield contrast-to-voltage impulse response functions for the LMC and the extracellular space. Taking the ratio of the contrast-to-voltage impulse response and the current-to-voltage impulse response in the Fourier space and performing inverse Fourier transform yields a contrast-to-current impulse response-a measure of effective injected current into the recording site as a response to unit contrast. This is seen to be roughly equal and opposite as seen in Figure 8 of Kashef and de Ruyter (2026) [2]. The definitions in Equations 11 and 12 make the assumption that the currents represented by these equal and opposite contrast-to-current impulse response functions are indeed the same Cl^−^ current. We note here that in Equation 9, a part of the chloride current is independent of the LMC transmembrane voltage. This represents the reversal potential induced current, and is the main part of the chlo-ride currents in our simulation. Noting from Equation 8 that *g*_Cl_ is event induced, we can define a pair of event to (reversal potential induced) voltage impulse response functions as

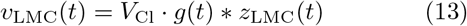

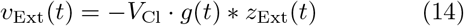

These responses are shown in Figure 3. While our results incorporate the corrections from the transmembrane potential as well, these potentials are quite informative, since they show us the basic voltages that are added to future voltages when a vesicle exocytosis event occurs. Each release event causes the extracellular potential to go up, causing the photoreceptor transmembrane potential to drop. From Figure 1, we see that this lowers the vesicle release rate and depresses the future release of vesicles-something we believe sets up regularization of vesicles. Also interesting is how the ratio of the amplitudes of these potentials reflect the ratio of the amplitudes of the respective contrast impulse responses (see Figures 2 and 10), even though they are constructed from our current to voltage impulse responses.

**FIG. 3.**
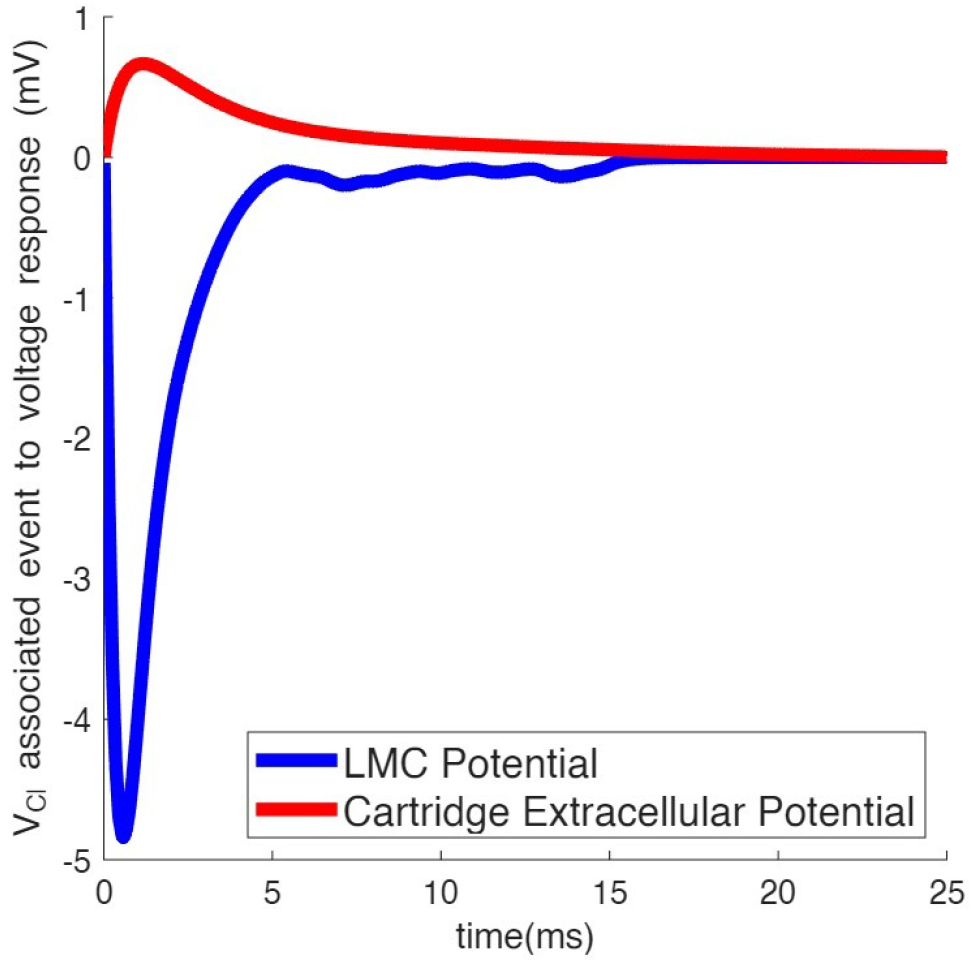
Reversal potential associated voltage responses of the simulated LMC and the extracellular space to a single release event.

**FIG. 4.**
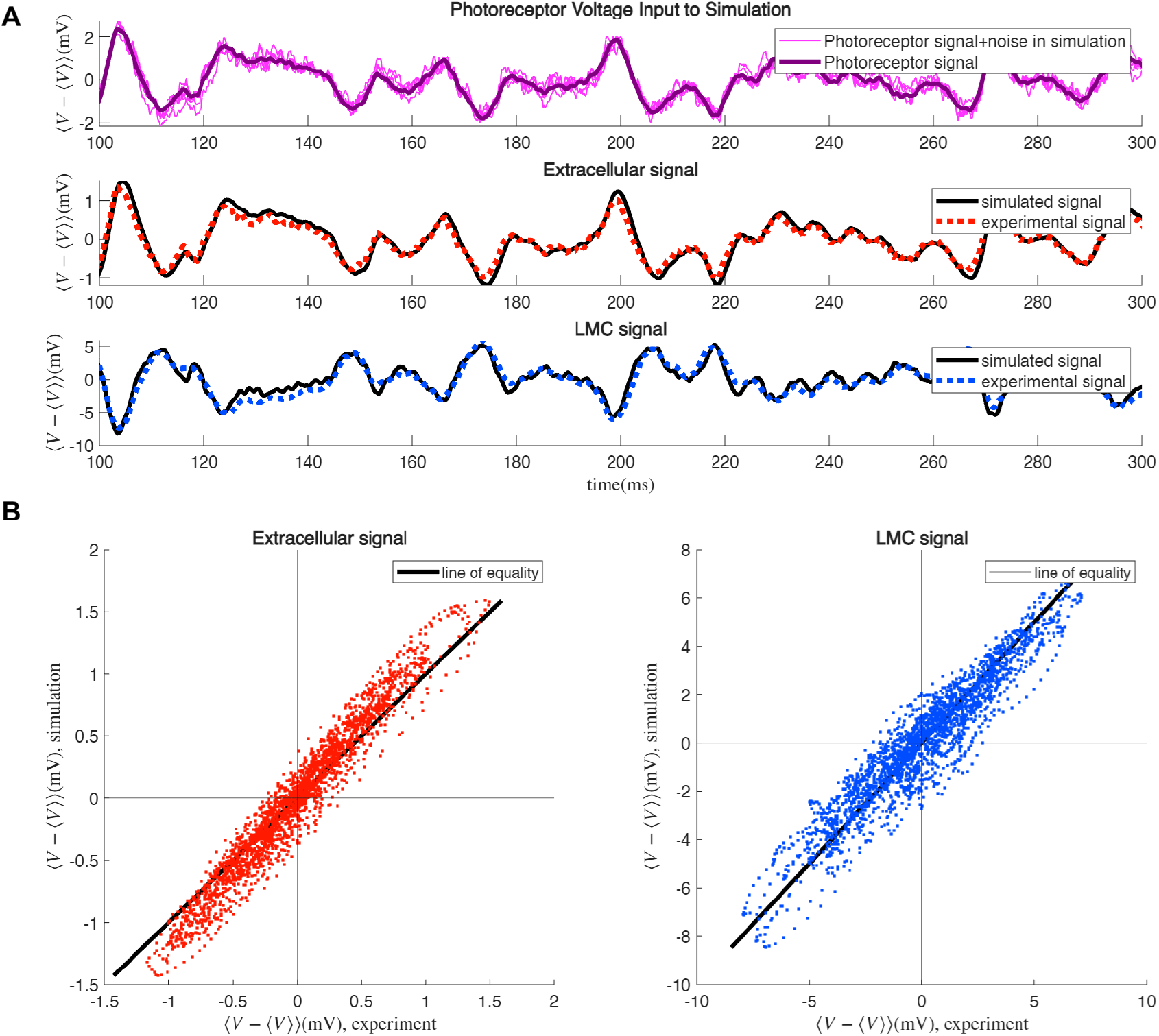
**A:** AC component of photoreceptor input signals to simulation (experimentally derived, with modified noise), with the trial averaged signal overlaid (*top*); extracellular output signal averaged over 200 simulations in black, with experimental signal shown in red dotted line (*middle*); and LMC output signal averaged over 200 simulations in black, with experimental signal shown in blue dotted line (*bottom*).**B:** AC component of signal in simulation versus the experimental counterpart for the extracellular (*left* ) and LMC (*right* ) voltages, overlaid on the line of equality.

### F. Summary of Parameters

In Table I, we summarize the fixed parameters of our computational model, their values and their sources. The free parameters were adjusted such that the signal and noise of the output extracellular and LMC potentials reasonably match the experimental analogs (i.e. extracellular and LMC potentials measured in the same cartridge as the photoreceptor potential used to drive the simulation). Table II shows us the list of the free parameters from our computational model, and three different combinations of those free parameters which worked to this end.

**TABLE I.** Fixed parameters in our simulations, each with a short description, the source of the value, and the value listed.

| Parameter | Description | Source | Value |
|---|---|---|---|
| *b* | Slope factor of the logistic function which determines the local time vesicle release rate as a function of the photoreceptor transmembrane voltage | LMC-photoreceptor signal amplitude relations, Laughlin et al. (1987)[11]; further justified by ion channel physics and known properties of voltage gated Ca<sup>2+</sup> channels[12]. | 0.56mv<sup>−1</sup> |
| *G*<sub>Cl</sub> | Average conductance of an open Cl<sup>−</sup> channel in postsynaptic LMC membrane. | Result of inside-out patch study of spontaneous opening of Cl<sup>−</sup> channels by Skingsley et al. (1995)[13] | 60pS |
| *τ*<sub>open</sub> | Time constant in distribution of Cl<sup>−</sup> ion channel opening periods in the postsynaptic LMC membrane. | Result of the same Skingsley study that gives us *G*<sub>Cl</sub>[13] | 0.5ms |
| *V*<sub>Cl</sub> | Chloride reversal potential of LMC membrane | Using the Nernst equation | −42.3mV |

**TABLE II.** Free parameters in our simulations, each with a short description and the choice of values in three different simulations.

| Parameter | Description | Simulation 1 Value | Simulation 2 Value | Simulation 3 Value |
|---|---|---|---|---|
| *R*<sub>max</sub> | The saturation point of the rate of vesicle release. | 1 × 10<sup>5</sup>s<sup>−1</sup> | 2.5 × 10<sup>6</sup>s<sup>−1</sup> | 2.5 × 10<sup>6</sup>s<sup>−1</sup> |
| *V*<sub>act</sub> | The ”activation” voltage, i.e. the value of transmembrane voltage for which the rate is half of *R*<sub>max</sub>. | −22mV | −19mV | −17mV |
| *N*<sub>Cl</sub> | The number of Cl<sup>−</sup> channels which open up as a result of a single synaptic vesicle release. | 400 | 400 | 100 |

## II. RESULTS

In all cases in Table II, the actual average vesicle release rate is somewhere between 4400 to 8000 per second, or between 3 to 6 per second per active zone, around a hundredfold lower than the measured maximum effective Poisson rates. Here, we focus on the results of simulation 1.

The top panel of Figure 4A shows the photoreceptor signals used to drive the simulation, overlaid on top of each other to represent the variability. The two lower panels show the simulation derived and experimentally measured LMC and extracellular potentials. The experimental signals were all collected from the same lamina cartridge. Figure 5 shows the contrast impulse response function (calculated using the original light contrast function used to collect the experimental data) for the simulations with the experimental contrast impulse function overlaid. The model works well to capture the shape of the signal in time, with contrast impulse responses very similar to experimental ones (with perhaps some sharper features).

**FIG. 5.**
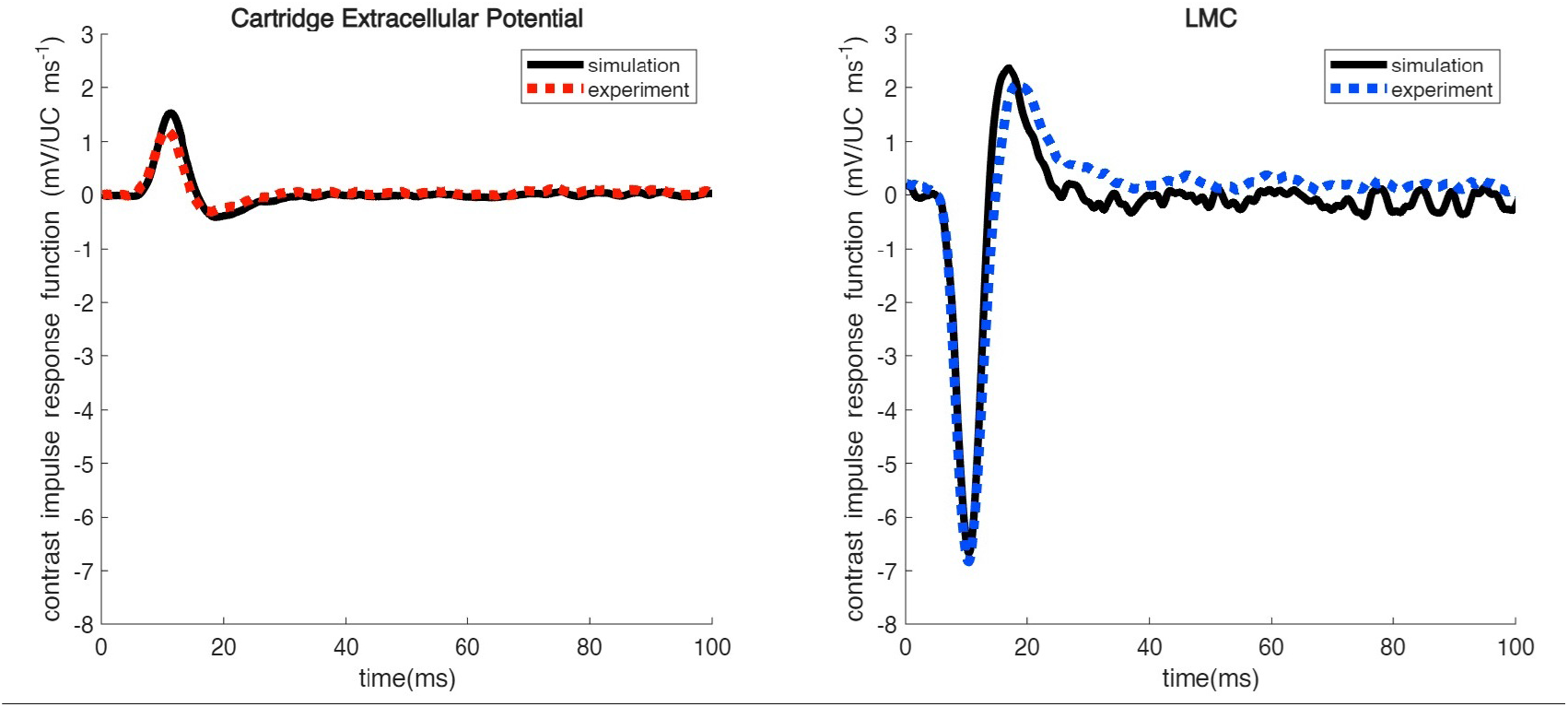
Contrast impulse response function of outputs from Simulation 1 in black, with corresponding experimental analogs overlaid in dotted line, for the cartridge extracellular potential (left) and the LMC (right).

Figures 6 and 7 show the computed contrast transfer function, noise power spectral density and effective Poisson rates for the extracellular potentials and LMCs in our simulations with experimental data overlaid. On the *R*_eff_ plot, the actual average rate of vesicle release events (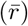) is also shown, showing around a two decade drop from the effective Poisson rate. This suggests that vesicle re-lease events in the simulation do not occur independently of each other, even though the simulation determines the number of events at a timestep only based on a local rate, and not based on previous event times. This is because when a release event occurs in the simulation, we not only add a negative vesicle induced potential to future timesteps for the LMC, but a positive vesicle induced potential for the extracellular space as well. Since, in our model, the rate of release depends on the instan-taneous *difference* in the photoreceptor and extracellu-lar potential, a release event *does* affect the probability of future events via negative feedback. The local drop in rate, in turn, lowers the number of potential generating events, and causes the rate to go back up, leading to a cycle of rate drops and increases, or regularization of vesicles. A scenario to this effect is explored in Kashef and de Ruyter (2026) [2] as a toy model. Plotting the event-triggered average of future events (this is also the autocorrelation function of the delta function train *ρ*(*t*) = Σ _*i* ∈events_ *δ*(*t − t*_*i*_) scaled by a factor of 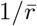 ) we find that indeed, event times are not uncorrelated, and have a reduced likelihood of occurring close to one another (although, because of the delay in response, the minimum of the autocorrelation occurs not right after *t* = 0, but slightly later, see Figure 8).

**FIG. 6.**
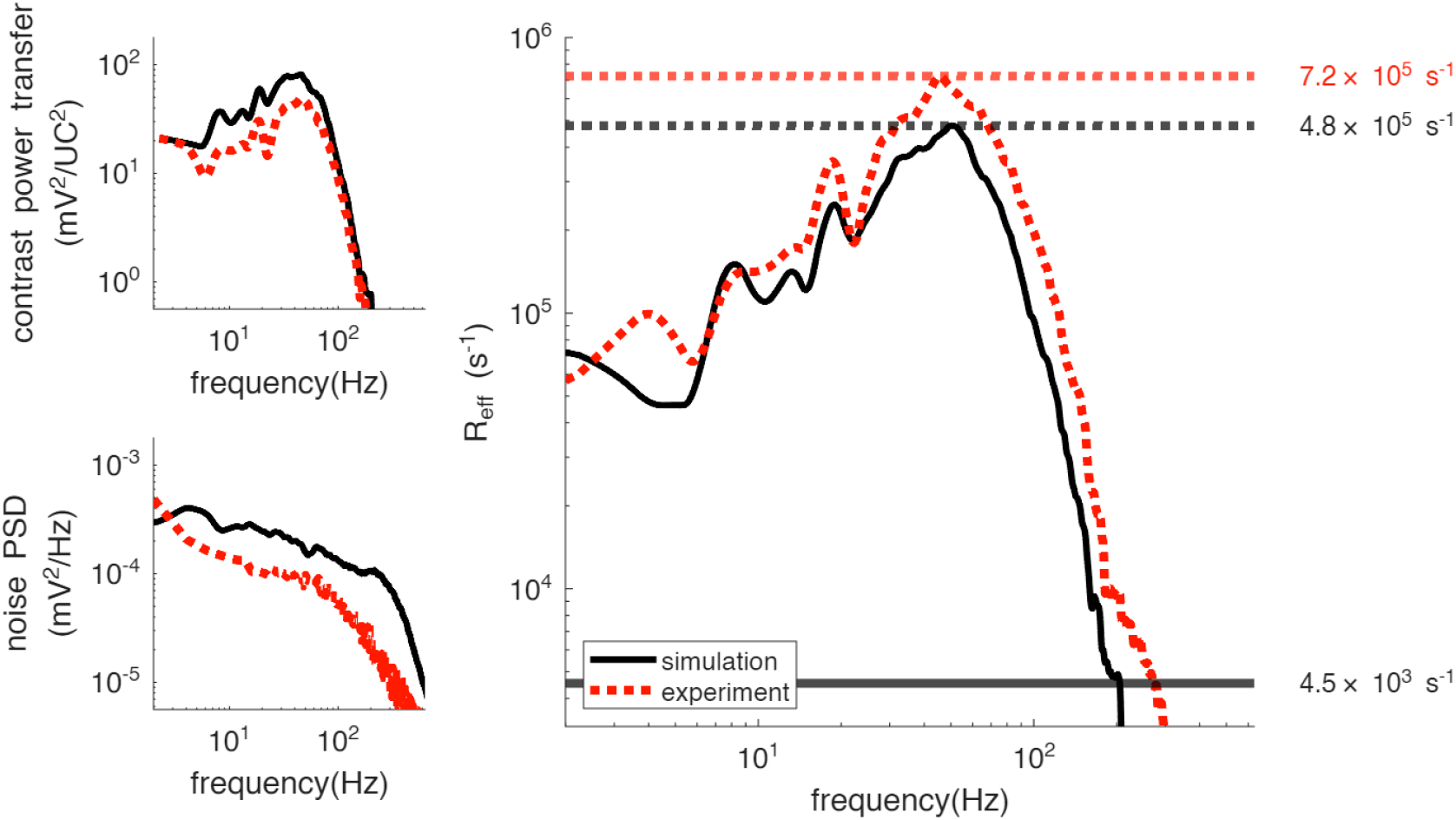
The contrast transfer function, noise power spectral density and the subsequent effective Poisson rates (in black) for the extracellular potential with the experimental analogs overlaid (in red dotted line). The maximum calculated effective Poisson rate is shown in a horizontal black dotted line for the simulation, and a horizontal red dotted line for the experiment, with the actual average rate from the simulation shown in a horizontal solid line.

**FIG. 7.**
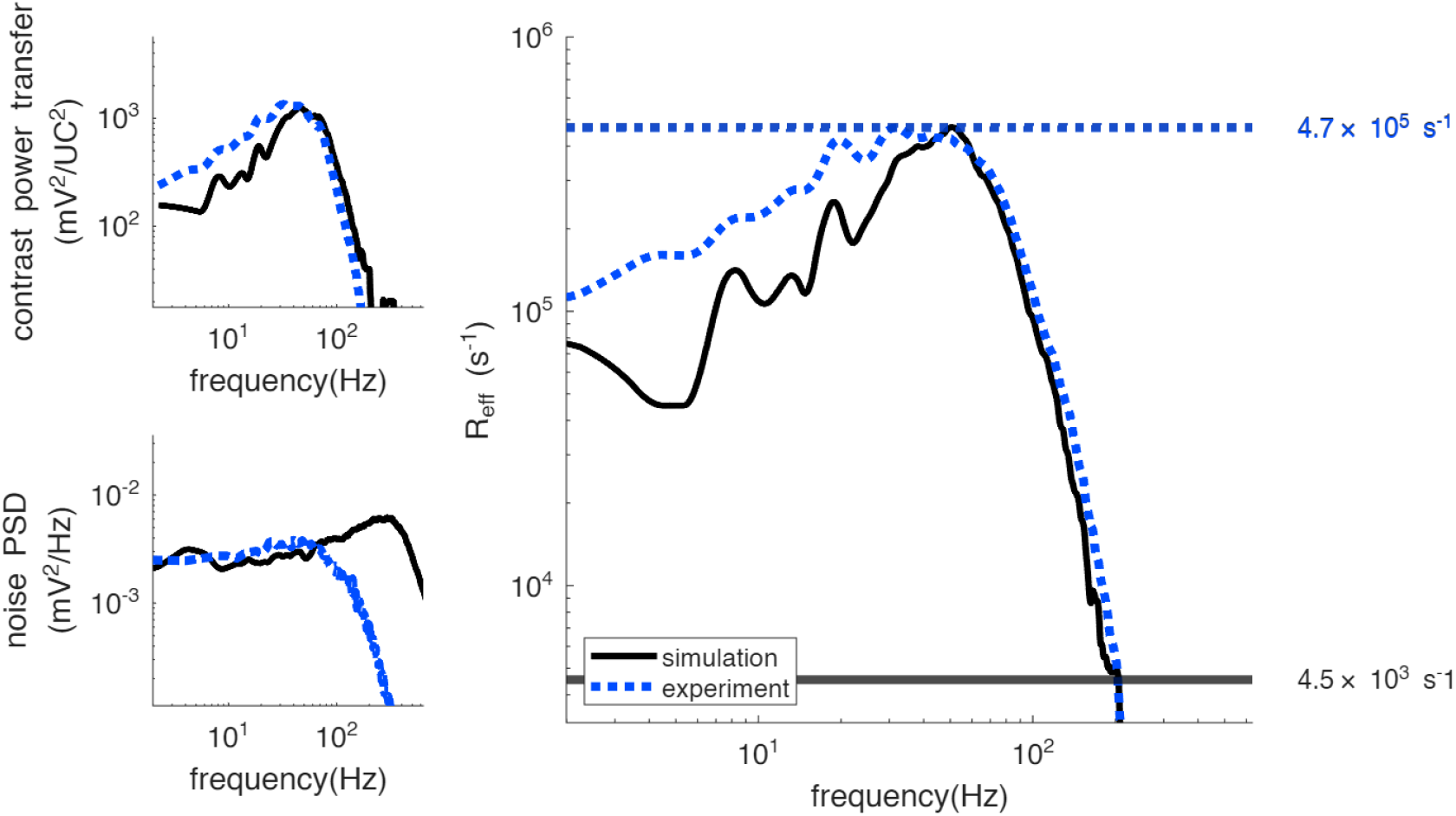
The contrast transfer function, noise power spectral density and the subsequent effective Poisson rates (in black) for the LMC potential with the experimental analogs overlaid (in red dotted line). The maximum calculated effective Poisson rate is shown in a horizontal black dotted line for the simulation, and a horizontal red dotted line for the experiment, with the actual average rate from the simulation shown in a horizontal solid line.

**FIG. 8.**
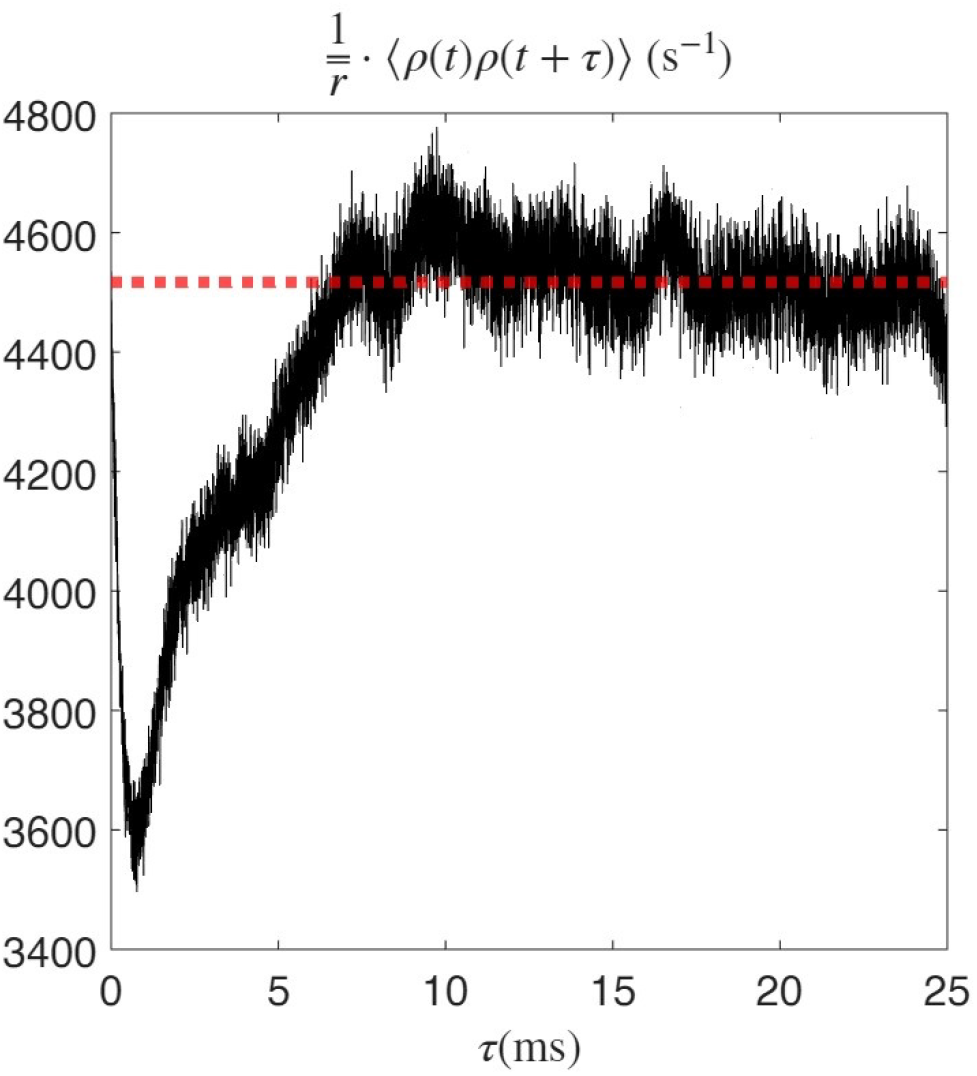
The autocorrelation function of the event delta function train in our simulation, normalized by the average rate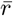 to give us the “vesicle triggered average” of future vesicle release events. The red dotted line shows the average rate of events.

## III. DISCUSSION

Since our picture of synaptic activity in the lamina involves feedback that affects future events, let us look at a simple tractable model that mimics this. As before, we start out with a rate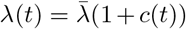 . We introduce a second time series *r*(*t*), which denotes the *actual* rate that the process follows (instead of *λ*(*t*)). We also define two variables:

- *r*_drop_, which determines a fixed amount that *r*(*t*) drops immediately in response to a release event
- *T*_relax_, which determines the time it takes for this drop in the rate to relax.

These in turn define a response function

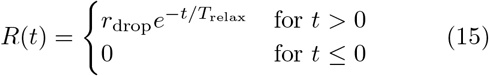

We initialize *r*(*t*) as *λ*(*t*), and every time an event occurs at some *t* = *t*_*n*_, we subtract *R*(*t* −*t*_*n*_) from the current *r*(*t*) during time-stepping. Thus, we have

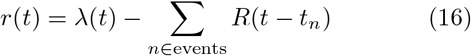

Combining the definitions in Equation 10 and 11, we infer that the system follows the following differential equation:

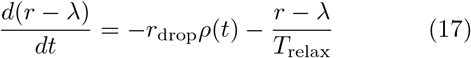

In the long run, the ensemble average of *ρ*(*t*) is just *r*(*t*). Applying ensemble averaging (which is valid for linear systems) and rearranging, we have

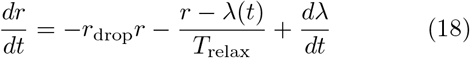

solving which gives us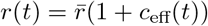, where

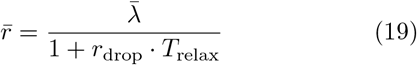

and

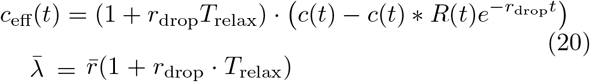

We set *T*_relax_ = 5ms in a numerical simulation, and try different values of *r*_drop_. In order to achieve actual average rates of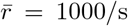, we have to set ·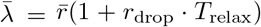 in simulations. The theoretical contrast transfer functions are given by 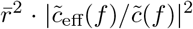 . The computed contrast transfer function, noise power spectral density and effective Poisson rates are shown in Figure 9. We see that with increasing regularization, the effective Poisson rate is given a boost from an increase in the contrast transfer function in the high end of the spec-trum *and* a reduction in the noise power spectral density on the low end of the spectrum. The former is tied to the fact that when event timing becomes more regular, the underlying rate can be inferred from fewer samples, particularly when observations are made over time windows longer than the typical inter-event interval. Figure 10 shows the *ρ*(*t*) that follows an event, averaged over all events. The value of this event-triggered average, of course, converges to the average rate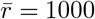 in the long run.

**FIG. 9.**
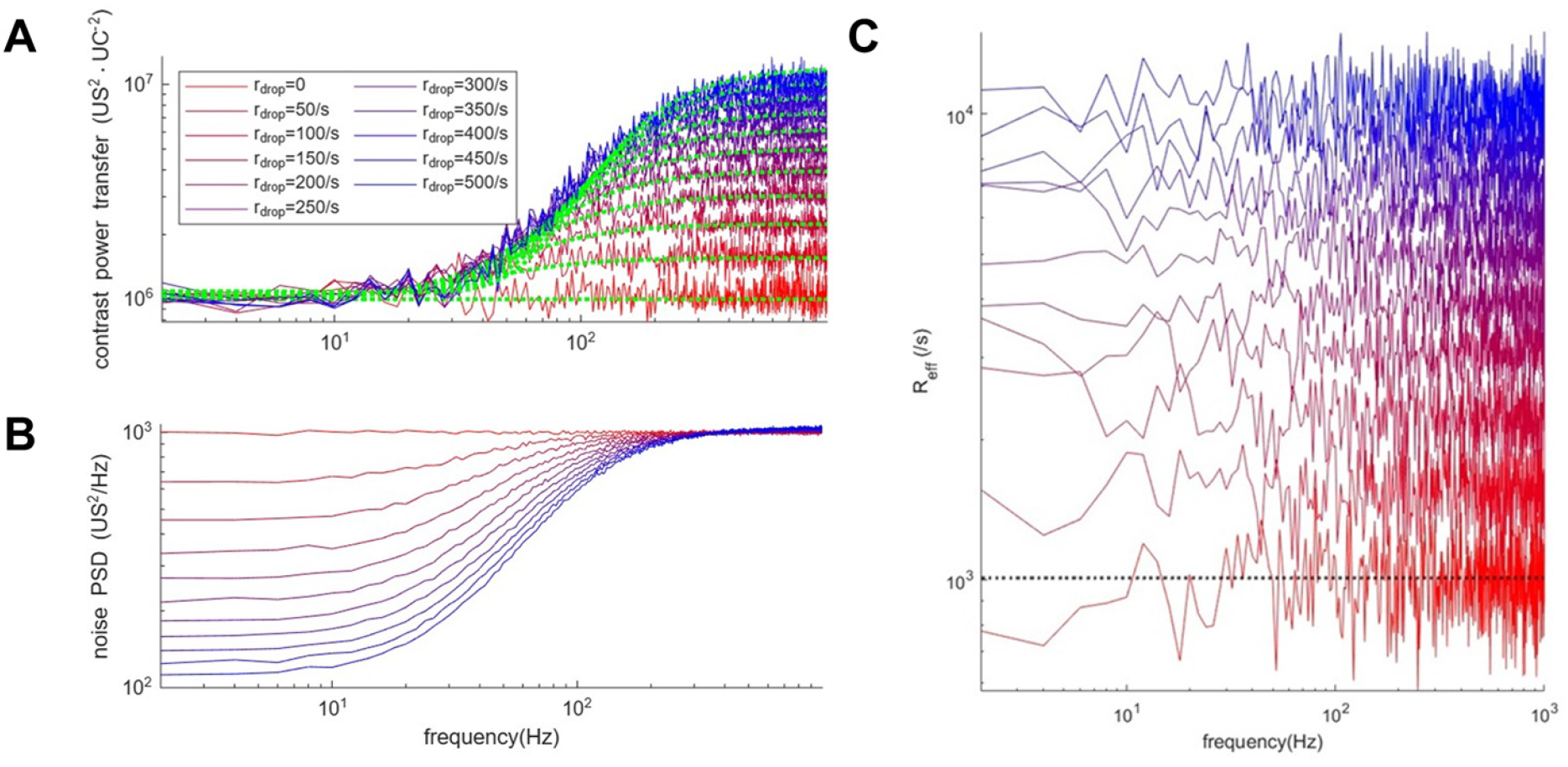
The contrast transfer function (**A**.), noise power spectral density (**B**.) and effective Poisson rate (**C**.) for processes with modulated “Poisson” rates where the current rate is determined by the occurrence of previous events. The mean event rate is chosen to be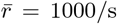, and the “contrast” is derived from the contrast function *c*(*t*) defining the raw underlying 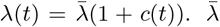 is modified in each case to maintain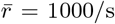 . The black dotted line represents where the effective Poisson rate equals the mean rate. The green dotted lines show the plots of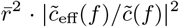, with *c*_eff_(*f* ) defined as the Fourier transform of the expression in Equation 20.

**FIG. 10.**
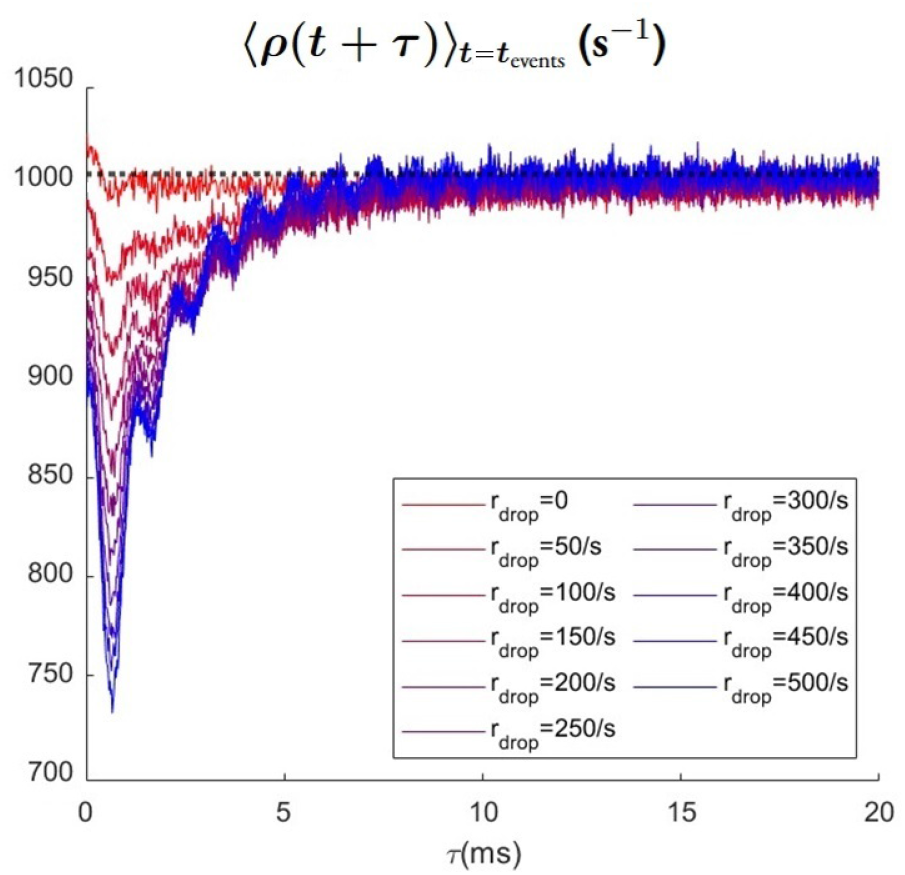
Event-triggered average of the Delta function train of future events for different values of *r*_drop_ and *T*_relax_ = 5ms.

A well-studied phenomenon in neurobiology where the synaptic activity at a given time immediately inhibits further synaptic activity is short-term synaptic depression in spiking neurons-the result of depletion of vesicles from synaptic activity [17]. Goldman (2004) frames the question of transmission efficiency in spiking neurons as bits per vesicle, and shows the improvement of this efficiency in certain conditions when presynaptic spikes fail to elicit vesicle release.[18] Here, the entropy in vesicle release times is defined as the “signal” portion of the contribution to transmitted information, and the entropy of the presynaptic spike times conditional on vesicle times as the subtracted “noise” portion. It is found, particularly in the case of depressing synapses (modeled as vesicles lowering the probability of release by an exponen-tial kernel, almost exactly as we do here) that the noise contribution per vesicle is *lower*, and thus information carried per vesicle is *higher* the stronger the depressive effects (i.e. higher *r*_drop_, *T*_relax_).

On top of the low frequency noise reduction that comes from increased predictability of event timings, the above analysis shows that a structured mechanism producing regularization (not unlike the mechanism explored in our model of the synapse) can amplify contrast power transfer, thereby contributing to an increase in the maximum effective Poisson rate. By defining *r*(*t*) as the ensemble-averaged rate across simulation trials, one can now introduce an “effective” contrast,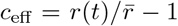, which differs from the contrast signal used to modulate light intensity in the experiments (see Figure 11). The lower panel of Figure 11 shows that the peak power of this effective contrast is enhanced by approximately an order of magnitude relative to the flat-spectrum light contrast. Consistent with the discussion above, regularization is expected to suppress the noise power spectral density, especially at low frequencies. Figure 12 illustrates the noise power spectra obtained by convolving event trains from Simulation 1 with the average event-evoked voltage response (both extracellularly and in the LMC) over 200 trials, and compares this with an equivalent convolution using statistically independent events generated from 200 repeated realizations of the modulated rate *r*(*t*). The results show an approximately tenfold reduction in noise power in the regularized simulation relative to the Poissonian case. Combined with the increased peak power in *c*_eff_, this reduction accounts for the observed hundredfold increase in the effective Poisson rate.

**FIG. 11.**
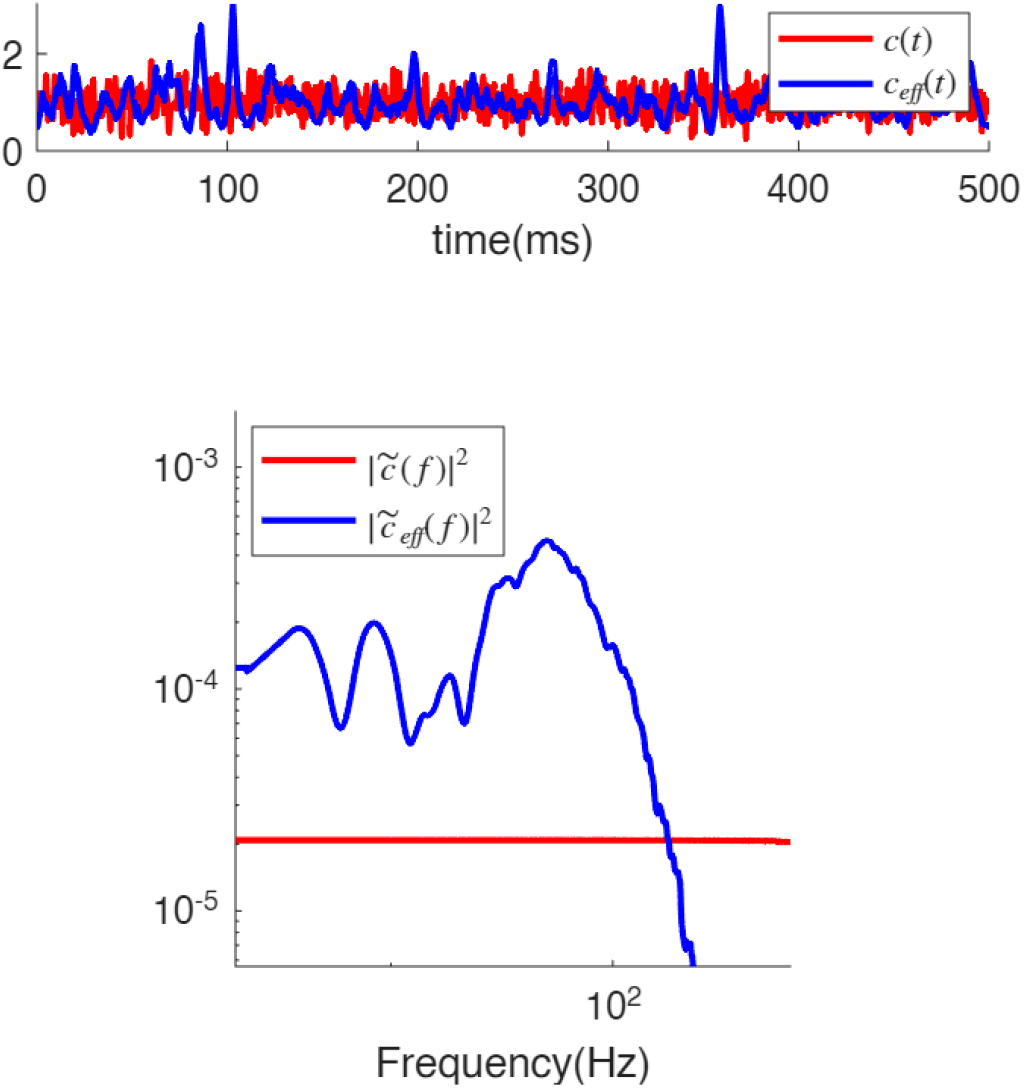
Time series (top) and power spectra (bottom) of the LED contrast in experiments (red) and the “effective” contrast from simulations (blue).

**FIG. 12.**
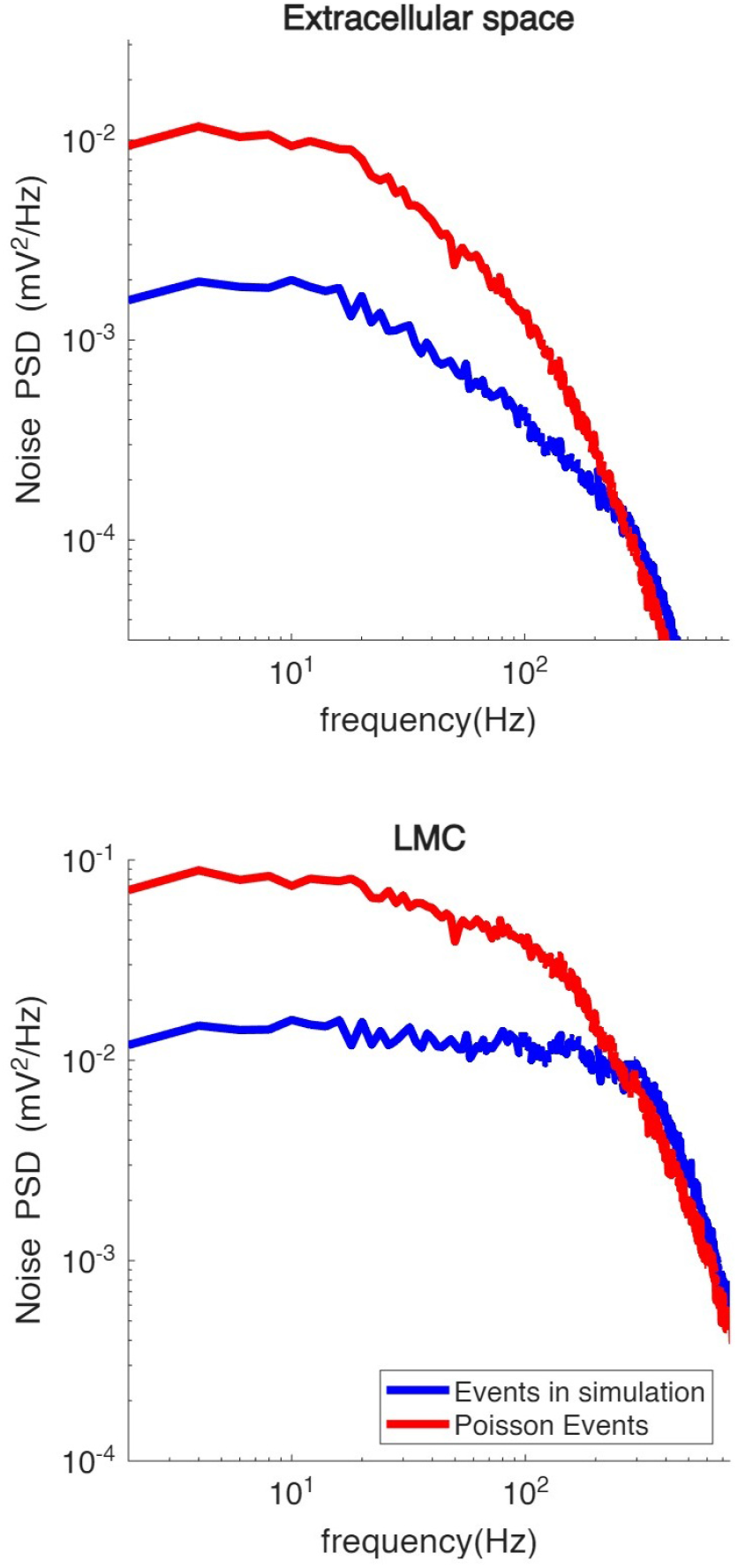
Comparison of the noise power spectral density for the original simulation and a Poisson event train with the same mean rate as simulation 1 convolved with the average voltage response per event for the cartridge extracellular potential (top panel) and the LMC (bottom panel).

How does each free parameter in Table II influence the simulation outcomes? Increasing *R*_max_ generally raises the mean event rate and enhances the sensitivity of the local release rate to changes in the photoreceptor transmembrane voltage (note that the slope of the rate function at its point of maximal susceptibility is *R*_max_*b/*4). Values of *R*_max_ that yield good agreement with measured contrast transfer and noise levels fall in the range of 10^5^ to *>* 10^6^ per second, depending on the remaining parameters. When divided across 1320 active zones, this corresponds to ∼75 to ∼2000 vesicles per second per active zone, when all Ca^2+^ channels are open and there are no other limiting factors. Increasing *V*_*act*_ tends to reduce the release rate by shifting the rate function in Equation 6 to higher voltages, such that the same photoreceptor membrane potential produces a lower release probability. In general, higher release rates reduce noise and improve the effective Poisson rate; however, event regularization also plays an important role. This effect is controlled by the final free parameter, *N*_Cl_, which sets the strength of extracellular feedback per vesicle by scaling the conductance associated with individual release events (Equation 7). For comparison, classical studies of large synapses (e.g., Katz-type preparations) estimate that a single synaptic quantum opens roughly 100–1000 postsynaptic channels [19]. The values used here fall within the lower to intermediate portion of this range. The value of *N*_Cl_ can be made smaller than this, generally raising the actual rate of vesicle release, but the numbers presented here lead to reasonable matches with the observed noise levels.

The picture in Figure 2 is highly dependent on the suggestion from the results of Kashef and de Ruyter (2026), that the current into the extracellular space is directly related to the chloride entering an LMC. Using the measurements of van Hateren and Laugh-lin (1989) [20], we can estimate the total axonal ca-ble resistance. Taking their reported membrane resistivity of 5 ×10^6^MΩ ·µm^2^ and axial resistance per unit length of 0.8MΩ*/*µm^2^, together with cable dimensions of *l* ≈ 425µm and *r* ≈ 1.35µm, yields a total DC resis-tance of approximately 1.5GΩ (calculated numerically). Since this cable impedance is so high, our model cannotrely on it to generate the observed current-to-voltage impulse response *z*_LMC_. We propose a workaround here, incorporating the distributed property of the extracellular impedance, given that the extracellular space is a tortuous one, and the recording site is separated both functionally and physically from the site of actual synaptic activity by a maze of membranes. Figure 13 illustrates a simplified full-circuit representation of the system, including pathways to ground. While a more detailed model seems desirable, this is the most detail we can work in, given the data and parameters available to us, in order to demonstrate the possibility of generating observed voltage-to-current responses without excessive speculation.

**FIG. 13.**
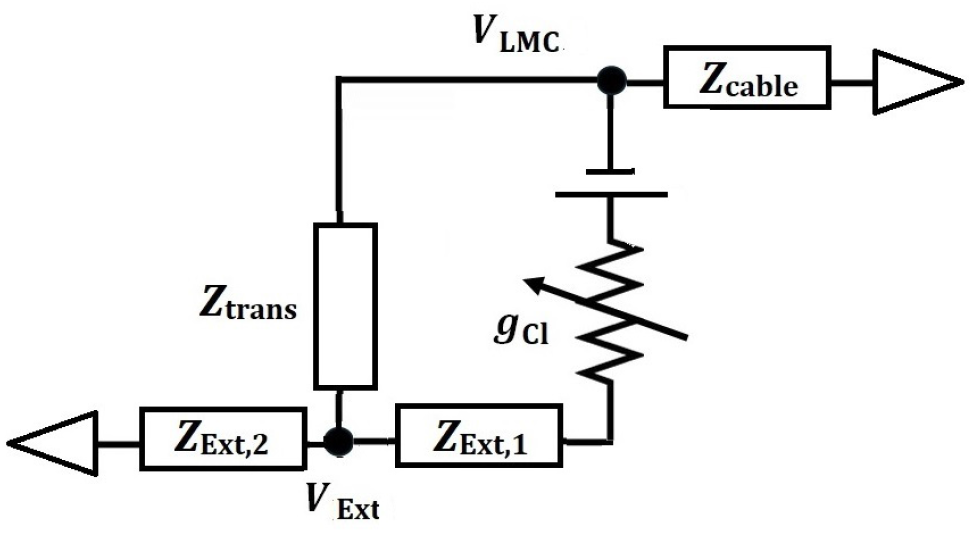
A representation of the cartridge circuit showing all known and potential impedance components, excluding any connections to the photoreceptors.

In this framework, *V*_Ext_ denotes the extracellular potential at the recording site, with the resistance to ground treated as *distributed* due to the tortuous geometry of the synaptic region [8, 21]. The impedance between the synaptic cleft and the extracellular recording electrode is denoted *Z*_Ext,1_, corresponding to the transmembrane pathway from the LMC to the recording site. The frequency-dependent cable impedance, *Z*_cable_is computed directly from the parameters reported by van Hateren and Laughlin.

Because we observe the response at only two points in the circuit following current injection, and have independent knowledge of just *Z*_cable_, it is not possible to uniquely determine *Z*_trans_, *Z*_Ext,1_ and *Z*_Ext,2_ simultaneously. Instead, we examine two limiting but plausible configurations. Calculations of chloride-mediated responses are therefore carried out under two distinct as-sumptions:

- **Case 1:** There is no appreciable conductance between the extracellular recording site and the LMC (i.e.,*Y*_trans_ = 1*/Z*_trans_ is set to zero in the fit). In this scenario, the recording site is separated from the synaptic cleft by *Z*_Ext,1_ and from ground by *Z*_Ext,2_.
- **Case 2:** There is a finite conductance between the extracellular recording site and the LMC, while *Z*_Ext,2_ separates the recording site from ground. The synaptic cleft and the recording site are assumed to be equipotential, and thus *Z*_Ext,1_ is set to zero in the fit.

In both cases, the free parameters are chosen such that current injection reproduces the experimentally observed voltage responses, with *Z*_cable_ held fixed. Figure 14 shows the resulting chloride-induced voltage transients for a single synaptic event in both the extracellular space and the LMC for Cases 1 and 2, alongside the kernels used in the simulations. In case 1, the inclusion of the cable permits a configuration in which voltage transients generated by chloride ion flux reproduce the responses used in our simulations. In contrast, in case 2, where *Z*_Ext,1_ = 0, the large cable impedance attenuates the voltage responses in both the extracellular space and the LMC. Notably, the resulting extracellular potential is too small to support a negative feedback mechanism capable of producing appreciable regularization. This indicates that a non-negligible *Z*_Ext,1_ is likely required to generate the degree of vesicle regularization observed in the simulations—that is, there must be sufficient electrical isolation between the recording site and the synaptic cleft. A more comprehensive treatment will be needed to disentangle the roles of the various anatomical components in shaping chloride-driven voltage signals, including contributions from photoreceptor membranes, which are not considered here.

**FIG. 14.**
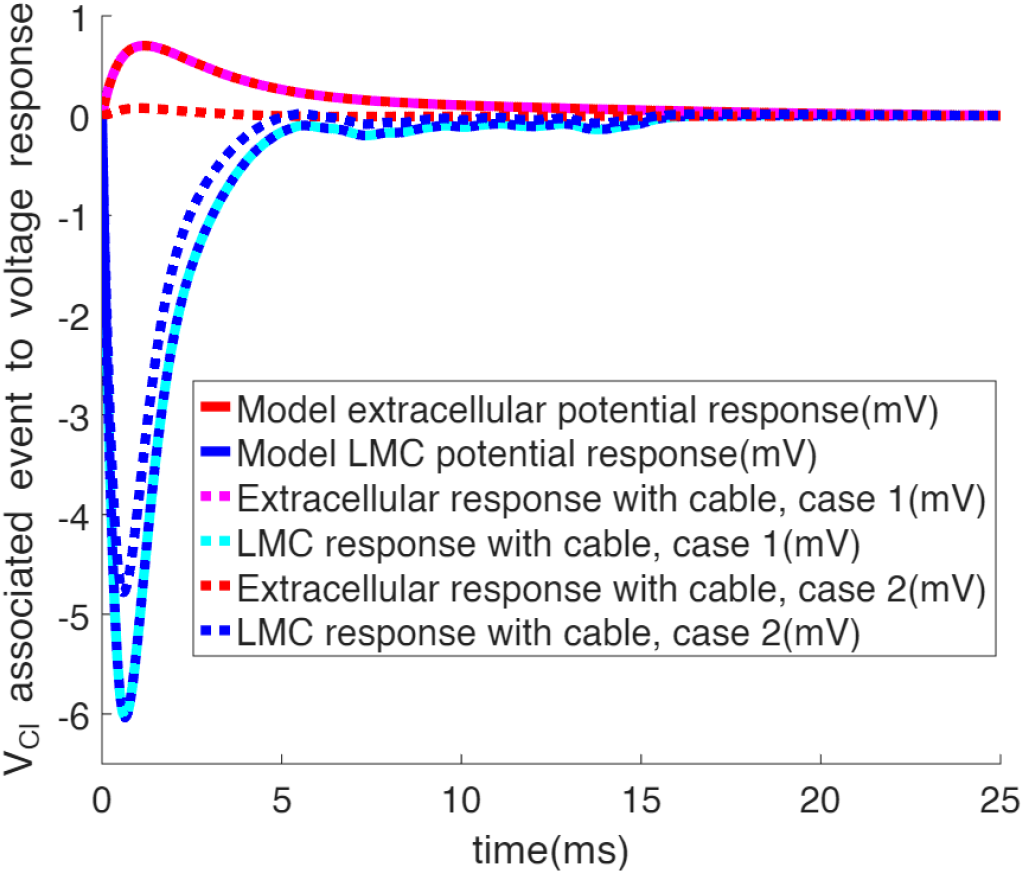
Voltage responses induced by a single chloride event in the synaptic extracellular space and in the LMC for Case 1 (dotted magenta and dotted cyan, respectively) and Case 2 (dotted red and dotted blue, respectively), shown alongside the extracellular and LMC kernels from Figure 3 (solid red and blue, respectively).

## IV. CONCLUSION

This study employs computational modeling to investigate visual information transmission in the lamina of *C. vicina*.The model builds on results from prior work [2, 11, 13], most notably the finding of Kashef and de Ruyter (2026) that the current entering an LMC is matched by an equal and opposite current leaving the cartridge extracellular space. With this constraint, the model contains three free parameters (Table II), which are varied to reproduce regimes in which the signal waveform and amplitude, as well as the noise power spectral density, are preserved. Experimentally measured effective Poisson rates from LMC voltage responses to flat-spectrum light contrast stimuli [2] are on the order of *>* 10^5^ per second, values that are difficult to reconcile with plausible vesicle release rates. In our simulations, however, the maximum effective Poisson rate exceeds the actual event rate by roughly two orders of magnitude, substantially reducing the required release rate to only a few events per second per active zone, rather than hundreds (Figures 6 and 7).

The model exhibits clear signatures of event regularization (Figure 8), arising from feedback potentials generated in the extracellular space that transiently suppress subsequent release events (Figure 3). As a consequence, vesicle release events are not statistically independent. Experimentally, the effective Poisson rates measured from extracellular cartridge potentials are of a similar magnitude to those observed in LMCs, suggesting a shared underlying mechanism shaping both signals-a relationship that is captured by the model.

We further examine how high maximum effective Poisson rates can emerge from discrete processes operating at comparatively low average rates through regularization. Regularization allows the maximum effective Poisson rate to exceed the mean driving rate of the process (Figure 9). Because each event consumes finite resources-not only metabolic energy for transmitter synthesis, but also structural resources such as axonal space required to sustain vesicle exocytosis-this amplification of effective rate relative to actual event rate enhances the efficiency of resource utilization.

Although this work suggests the presence of an adaptation that exploits the electrical properties of the LMCs and their surrounding environment to enhance the efficiency of information transmission, additional experiments are required to clarify the mechanisms operating at the synapse. Two outstanding puzzles remain. First, during identical light stimulation, the current measured in an LMC is equal and opposite to the extracellular current, despite the presence of two LMCs in the cartridge. Second, the high impedance of the LMC axon [20] would, in principle, strongly limit the ability to draw current from ground and thereby generate a feedback voltage of sufficient magnitude in the extracellular space. Both issues are resolved if the recording and current-injection site is electrically separated from the synaptic cleft by a substantial impedance (Figures 13 and 14). Further anatomical characterization and targeted electrophysiological measurements will be necessary to determine whether such electrical separation is indeed present and to what degree it shapes synaptic signaling.

